# The complement pattern recognition molecule CL-11 promotes invasion and injury of respiratory epithelial cells by SARS-CoV-2

**DOI:** 10.1101/2023.12.11.571109

**Authors:** Anastasia Polycarpou, Tara Wagner-Gamble, Roseanna Greenlaw, Lauren A. O’ Neill, Hataf Khan, Michael Malim, Marco Romano, Dorota Smolarek, Katie Doores, Russell Wallis, Linda S. Klavinskis, Steven Sacks

## Abstract

Collectin-11 is a soluble C-type lectin produced at epithelial surfaces to initiate pathogen elimination by complement. Given the respiratory epithelium is a source of CL-11 and downstream complement-pathway components, we investigated the potential of CL-11 to impact the pathogenicity of SARS-CoV-2. While the SARS-CoV-2 spike trimer could bind CL-11 and trigger complement activation followed by MAC formation, the virus was resistant to lysis. Surprisingly, virus production by infected respiratory epithelial cells was enhanced by CL-11 opsonisation of virus but this effect was fully inhibited by sugar-blockade of CL-11. Moreover, SARS-CoV-2 spike protein expressed at the bronchial epithelial cell surface was associated with increased CL-11 binding and MAC formation. We propose that SARS-CoV-2 pathogenicity is exacerbated both by resistance to complement and CL-11 driven respiratory cell invasion and injury at the portal of entry. Contrary to expectation, CL-11 blockade could offer a novel approach to limit the pathogenicity of SARS-CoV-2.

---

Complement is a major part of the innate immune system for detecting and eliminating invasive pathogens and signalling to the antigen-specific arm of the immune response^1^. That many successful pathogens resist or usurp complement is indicative of the potency of complement as an antimicrobial deterrent^2,3^.

Activation of the complement system is initiated by pattern recognition molecules (PRM). C1q is the main PRM for the classical pathway for detecting antigen-antibody complexes, leading to the assembly of C4 and C2 and subsequent cleavage of C3, the central complement component. Collectins and ficolins are PRMs for the lectin pathway, detecting carbohydrate signatures on the activating surface prior to assembly of C4 and C2 and cleavage of C3^4^. The alternative pathway is different because it has no specific PRM and appears to play a secondary role amplifying cleaved C3 deposited by the classical or the lectin pathway, using complement factors B and D^5^.

The discovery of collectin-11 (CL-11) has sparked new interest in the collectin family of PRMs^6^. Each CL-11 polypeptide includes a carbohydrate-recognition domain (CRD) and a collagen-like domain (CLD). These combine to form trimers and multiples of trimers with increased avidity for carbohydrate ligands typically fucosylated glycans and high mannose oligosaccharides^7,8^. Like the structurally similar mannan-binding lectin (MBL), CL-11 can interact with a wide variety of bacteria, viruses, and fungi^6,9^. But only CL-11 is synthesised in peripheral tissues^6,7^ whereas MBL is entirely produced by hepatocytes and present in the blood; and the large molecular size of MBL oligomers (400-700 kDa) may preclude extravasation. This compartmental separation has led to the idea that CL-11 acts as a guardian of the extravascular tissue space, particularly at epithelial surfaces where the secretion of CL-11 parallels that of other essential complement proteins^10^. In contrast, MBL may primarily serve the intravascular compartment to protect against haematogenous spread of infection.

Like other collectins, CL-11 physically combines with MBL-associated serine proteases (MASPs 1-3), of which MASP-2 is the main lectin pathway enzyme^11,12^. Upon contact of the lectin with an activating surface, MASP-2 mediates the cleavage of C4 and then C2 to form C4bC2b, i.e., the C3 convertase shared with the classical pathway. Subsequent cleavage of C3 and then C5 leads to the formation of C3a, C3b, C5a and C5b-9; these complement effectors promote inflammation, opsonisation, and formation of membrane attack complex (MAC, C5b-9) downstream of pathogen recognition.

Clinicopathological studies of SARS-CoV-2 infection have suggested a link between severe lung inflammation and detection of complement factors that specify the lectin pathway, such as MBL, MASP-2 and C3b^13^, coinciding with SARS-CoV-2 antigen detection^14,15^. The presence of microvascular thrombosis, also a feature of severe lung pathology in this condition^15^ is a hallmark of complement activation particularly by the lectin pathway proteases^16,17^. In addition, a genome-wide study has identified an association between COVID-19 disease severity and CL-11^18^. It is also important to emphasise that early structural modelling of the SARS-CoV-2 spike protein - instrumental for the receptor recognition and membrane fusion process of cell entry – predicted 12 putative sites with which with CL-11 could interact^19^.

Based on these clinical, genetic, and structural observations, we set out to investigate the ability of SARS-CoV-2 to interact with CL-11 and determine the impact of complement activation on the virus and on virus-infected cells at the portal of viral entry. Our study unequivocally demonstrates interaction of CL-11 with SARS-CoV-2 and specifically the spike protein. Moreover, it reveals a significant effect of bound CL-11 on viral infectivity of respiratory epithelial cells exacerbated by cell-autonomous activation of complement.

## Results

### 1: SARS-CoV-2 binds CL-11 by its carbohydrate recognition site

At the onset of the COVID-19 pandemic we predicted that the pattern recognition molecule, CL-11, which is produced at the respiratory epithelial surface^6,20^, would engage with SARS-CoV-2^19^. This assumption built on historical evidence of SARS-Co-V interacting with the related collectin, MBL^21^, and the modelled crystal structure of the spike protein of SARS-CoV-2, which is rich in glycan signatures that can be recognised by CL-11^19^. The starting point of our study was to investigate the ability of SARS-CoV-2 and its spike protein to interact with CL-11. We carried out binding assays with immobilised UV-inactivated SARS-CoV-2 and immobilised spike trimer to address the interaction with human recombinant CL-11 (rCL-11). Our results confirmed rCL-11 binding by SARS-CoV-2 and spike protein in proportion to the dose of the virus or spike protein (Fig. 1 a-d). It is important to note that CL-11 demonstrated equivalent binding to both the ancestral Wuhan spike protein and the later Omicron BA.1 variant spike protein (EC50 values of 7.6 (± 4.2) × 10^−9^ M and 6.0 (± 2.6) × 10^−9^ M respectively), Fig. 1d. Controls for non-specific binding of rCL-11 used the immobilised non-glycosylated protein BSA (Fig. 1a, b & c).

**Fig. 1.**
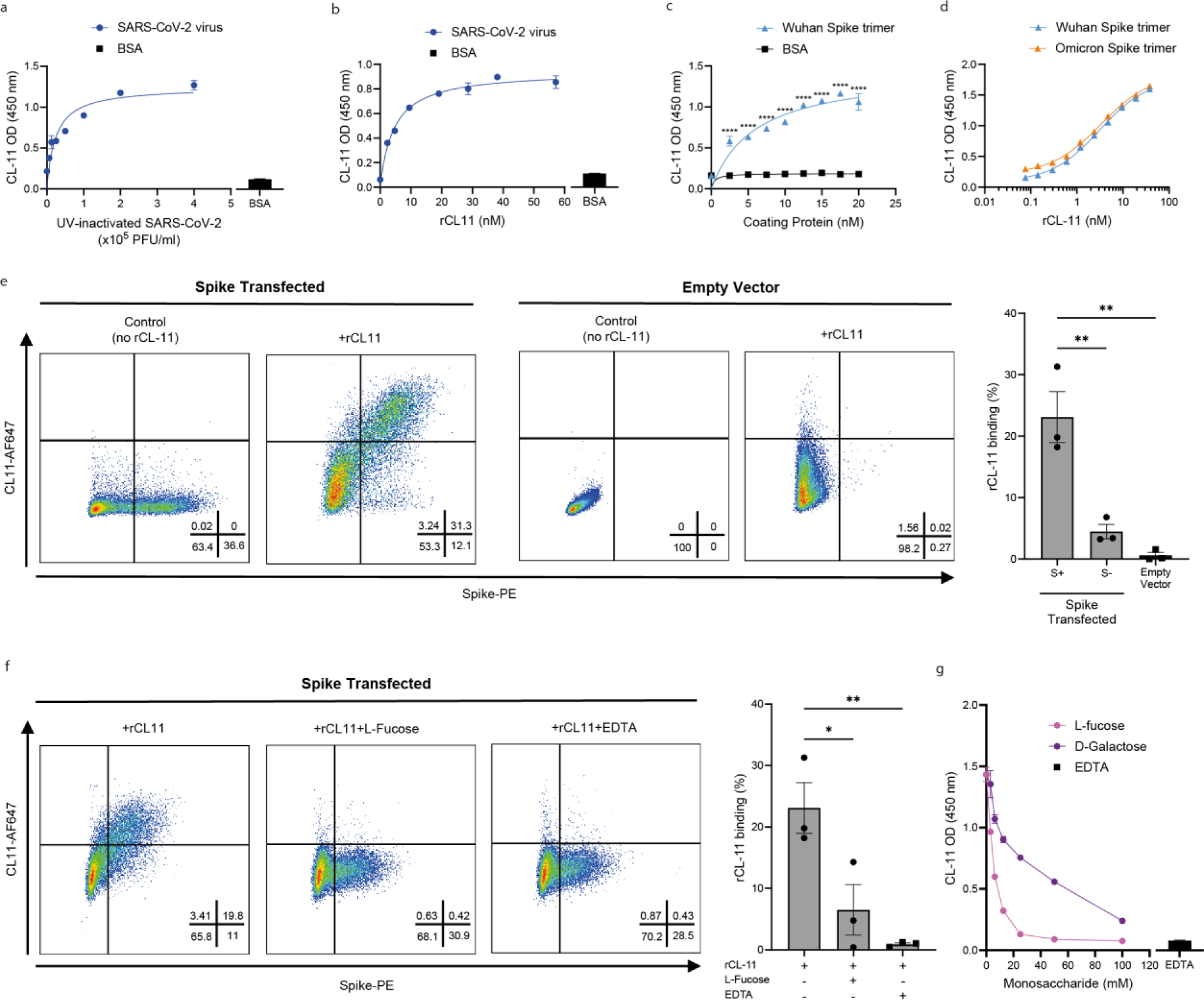
SARS-CoV-2 interacts with the carbohydrate binding domain of CL-11. **a-d**, Measurement of human rCL-11 bound to either UV-inactivated SARS-CoV-2, or recombinant trimeric spike protein or BSA immobilised on 96-well Maxisorp plates and detected by ELISA with specific antibody to human CL-11, HRP-conjugated secondary antibody and TMB. **a,** Quantification of bound rCL-11 following incubation of a fixed concentration (3µg/ml) with immobilised SARS-CoV-2 virions (6.25 ×10^4^– 4×10^5^ PFU equivalent /ml) or to BSA as a control protein. **b,** Quantification of rCL-11 bound to a fixed concentration of immobilised SARS-CoV-2 virions (2×10^5^ PFU/ml equivalent) or to BSA as a control protein. **c**, Quantification of bound rCL-11 following incubation of a fixed concentration (3µg/ml) with immobilised recombinant Wuhan spike trimer or to BSA over a concentration range. ****p< 0.0001 by two-way ANOVA with Šidák’s correction, error bars show S.E.M. **d,** Quantification of rCL-11, demonstrating binding (over a concentration range) to immobilised recombinant spike trimer Wuhan or Omicron (2µg/ml). **a-d,** Data representative of at least three independent experiments. **e,** Representative flow cytometry histograms and summary bar graph demonstrating rCL-11 bound to cell-surface expressed spike protein. Left two panels, spike-plasmid transfected HEK293T cells, and right two panels, empty-vector transfected HEK293T cells, treated with either PBS (control) or rCL-11 (25 µg/ml) prior to staining for cell surface CL-11 and spike protein. Bar graph summarises percent of cells with bound rCL-11 within the spike transfected cell population gated for cell surface spike protein (+ /-) or percent of cells with bound rCL-11 within the control (empty vector) transfected cell population from three independent experiments.**p < 0.01 by one-way ANOVA with Tukey’s multiple comparison, error bars show S.E.M. **f,** Representative flow cytometry histograms and summary bar graph demonstrating sugar blockade and Ca^2+^ dependence of rCL-11 binding to cell-surface expressed spike protein, for rCL-11 pre-treated with PBS, L-fucose (10mM) or EDTA (10mM) prior to incubation with spike transfected HEK293T cells. Bar graph summarises percent of cells with bound rCL-11 (mean ±S.E.M) from three independent experiments. *p< 0.05 and **p< 0.01 by one-way ANOVA with Tukey’s multiple comparison. **g.** Sugar specificity for CL-11 binding and inhibition to immobilised Wuhan spike trimer (2µg/ml), where rCL-11 (3 µg/ml final concentration) had been pre-treated with L-fucose or D-galactose (0-100 mM) to block carbohydrate recognition of CL-11, or with EDTA (10 mM final concentration) by ELISA to demonstrate Ca^2+^ binding-dependence.

We next assessed whether membrane-expressed spike protein would also bind CL-11. For this, we incubated spike-transfected HEK293T cells with a fixed amount of rCL-11. Flow cytometry demonstrated significant binding of rCL-11 to cells expressing cell surface spike compared to transfected cells negative for cell surface expressed spike or empty vector (EV), control transfected cells (Fig. 1e). These data provide strong evidence that CL-11 readily interacts with SARS-CoV-2 spike protein (as predicted from the crystal structure) when the protein is expressed on cells or immobilised on plastic.

Our experiments went on to address whether the interaction with spike protein involved the CRD of CL-11. In theory binding of CL-11 to a target surface could occur either through the CRD of CL-11 or its CLD. Binding by the CRD of C-type lectins such as CL-11 has two distinct properties: Ca^2+^-dependent binding to the target substrate; and sugar-inhibitable binding. A series of experiments using HEK293T-cell-expressed spike trimer or plate-immobilsed spike trimer as the binding target determined rCL-11 binding to be both EDTA and sugar-inhibitable (Fig. 1f,g). Thus, treatment with increasing concentrations of L-fucose to saturate the CRD of CL-11 typically inhibited rCL-11 binding to the target by >90%. LIkewise, the Ca^2+^ chelator EDTA inhibited the CL-11/spike interaction by a similar amount. Inhibition was less effective with D-galactose control (Fig. 1g), which has been shown to have significantly lower binding avidity for CL-11 than L-fucose^7^.

We concluded from these initial experiments that the interaction between the SARS-CoV-2 spike protein and CL-11 is mediated by the CRD of CL-11, an interaction which is potentially blocked in the presence high concentrations of L-fucose.

### 2: SARS-CoV-2 triggers complement activation

Since our data indicated rCL-11 present at a concentration equivalent to the level in normal human serum (NHS; 0.1-0.5μg/ml; 0.95-4.76 nM) was bound by SARS-CoV-2 (Fig. 1b,d), we examined how well SARS-CoV-2 could stimulate complement activation in NHS, where other essential complement components are present. In the first of two independent approaches, we measured the soluble complement activation products C3a and C5a released into the supernatants after incubating SARS-CoV-2 virions with NHS as the source of complement. The levels of C3a and C5a generated were significantly above those in control samples in the absence of added virus (Fig. 2a,b). The basal levels of C3a and C5a detected in these control assays are likely to reflect the presence of these products in NHS^22,23^ or may represent low-level complement activation on the plastic surface of the assay system.

**Fig. 2.**
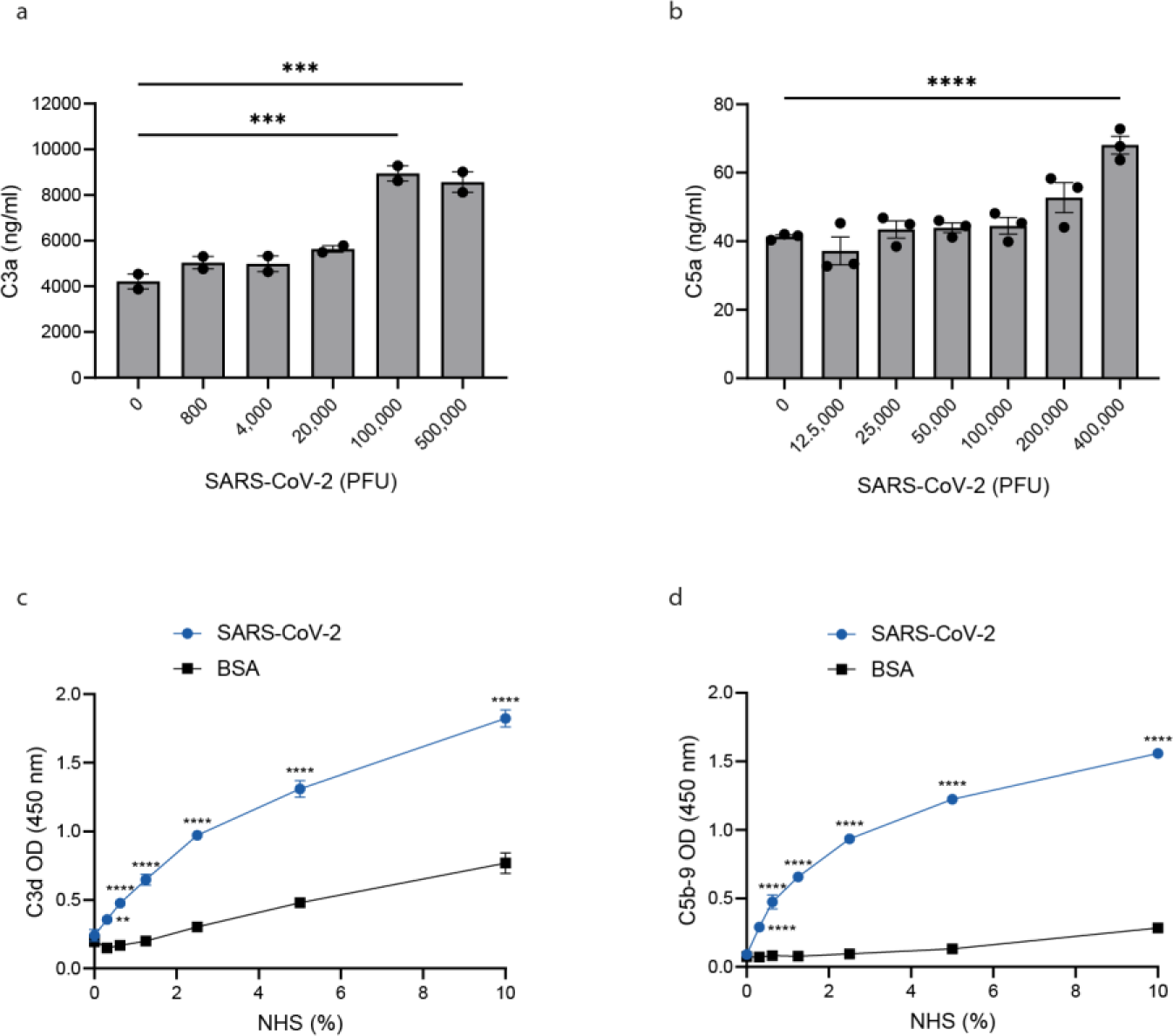
SARS-CoV-2 initiates complement activation. **a-b**, Quantification by ELISA of C3a and C5a generated in NHS after incubation (1:1 by volume) with live SARS-CoV-2 virions. Data shown is the mean ± S.E.M of triplicate biological samples for each virus concentration shown for **a**, C3a and **b,** C5a, and is representative of three and two independent experiments respectively. *P* values determined by one-way ANOVA with Dunnett’s multiple comparisons against no virus control. ***p< 0.001, ****p< 0.0001 **c-d,** Detection of C3d and C5b-9 deposited on SARS-CoV-2 virions. ELISA plates were coated with UV-inactivated virions (2 ×10^5^ PFU/ml equivalent) or BSA (as control), and incubated with NHS for 1 hr at 37°C. Deposited C3d (**c**) and C5b-9 (**d**) detected with specific antisera against C3d and C5b-9 followed by enzyme conjugated anti-species detection antibodies. Data shown is from triplicate samples and is representative of three independent experiments. ** p < 0.01 and **** p<0.0001 determined by two-way ANOVA with Šidák’s multiple comparison.

Using a second approach, we examined the deposition of C3d and C5b-9 on UV-inactivated SARS-CoV-2 virions immobilised on plastic wells, where C3d is the end metabolite of bound C3b deposited on an activating surface and C5b-9 is MAC formed after cleavage of C5. We detected significant increases of bound C3d and C5b-9, as compared with control measurements where immobilised BSA replaced the virus (Fig. 2c,d). The deposition of complement was dose-sensitive over a range of serum dilutions (0-10% NHS). These data indicate a low threshold for complement activation by SARS-CoV-2 under the assay conditions.

### 3: SARS-CoV-2 is resistant to complement-mediated lysis

Because COVID-19 can be a clinically aggressive disease, despite activation of the complement cascade (as confirmed in Fig. 2c,d), we questioned whether the virus could be resistant to complement-mediated lysis, as reported for some other viruses^24–26^. To test this we assessed the capacity of SARS-CoV-2 to resist virion lysis by NHS. We incubated SARS-CoV-2 virions with NHS and Ribonuclease A (RNase A) and then determined the ability of RNAse A to access the genomic RNA and degrade virions upon virolysis (Fig. 3a). In control studies where SARS-CoV-2 was treated with the detergent Triton-X instead of serum, as expected qPCR detected negligible genomic viral RNA (Fig. 3b). Conversely, the level of SARS-CoV-2 RNA was not statistically reduced when virus treated with NHS (1-10%) when cross-compared with untreated virus (Fig. 3b). To further probe the role of CL-11, we used NHS supplemented with rCL-11 in a final concentration (1-10µg/ml; 9.52nM-95.24nM) predetermined to give adequate binding with SARS-CoV-2 as in earlier experiments (Fig. 1a, b). However, the NHS enriched with CL-11 had no detectable effect on virolysis compared with non-enriched NHS (Fig 3c).

**Fig. 3.**
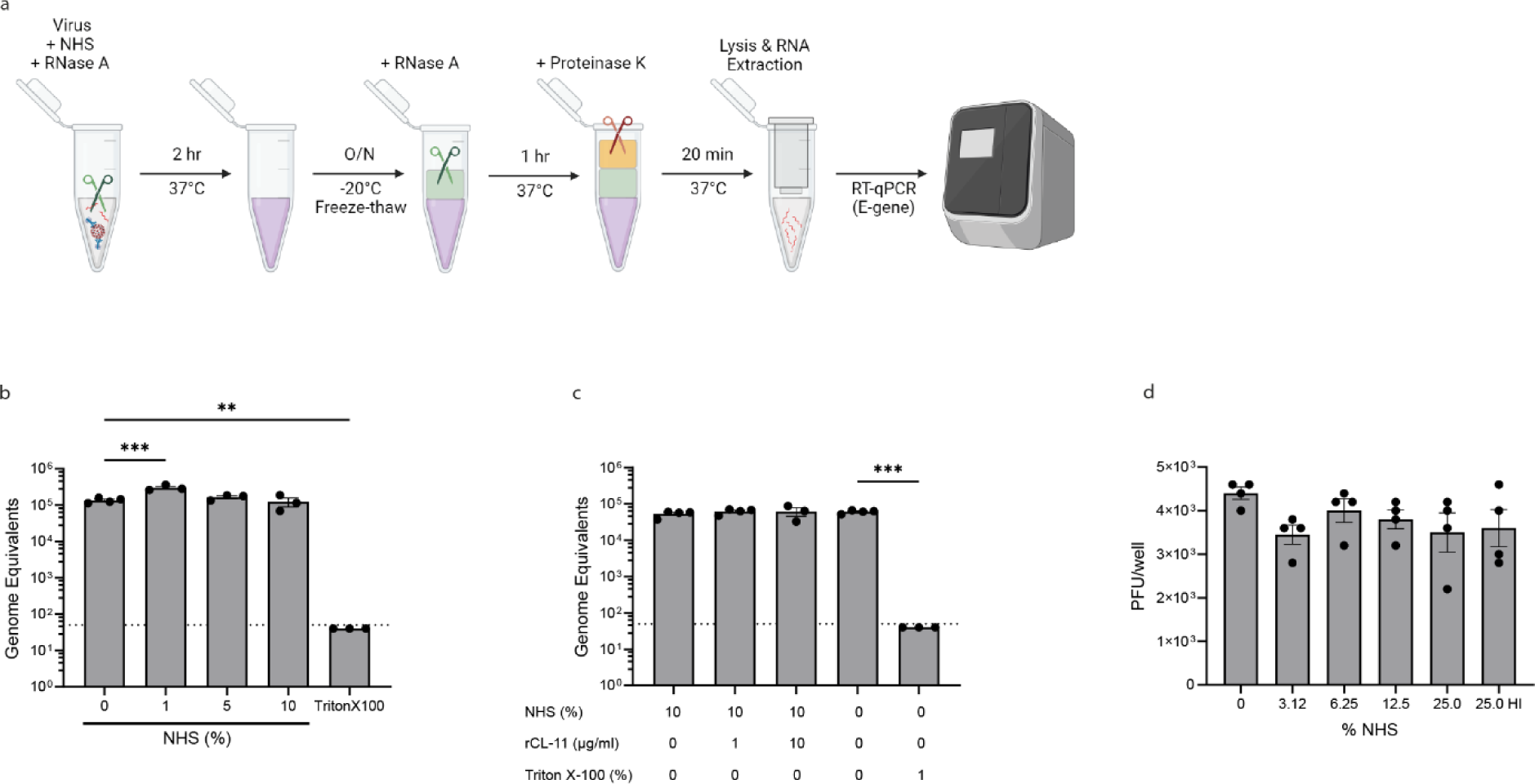
SARS CoV-2 virions are resistant to complement mediated lysis. **a**, Work flow to determine resistance of SARS-CoV-2 to complement by virion lysis assay, using pooled NHS (final concentration 0, 1, 5 and 10%) in the presence of SARS-CoV-2 virions and RNase A. Triton-X-100 (1% final concentration) was included as a control. Absolute viral RNA quantitation was determined by RT-qPCR and expressed as number of SARS-CoV-2 E genome copies. **b,** Virus lysis by pooled NHS (final concentration 0, 1, 5 and 10%) measured by RT-qPCR and expressed as number of SARS-CoV-2 E genome copies. Data represents the mean ± S.E.M of at least triplicate biological samples assayed in duplicate by RT-qPCR and is representative of three independent experiments. **c,** Resistance of SARS-CoV-2 virions to lysis by 10% NHS in the presence of rCL-11 (1.0 and 10.0 µg/ml final concentration) and RNase A. Triton-X-100 (1% final concentration) was included as a control. Absolute viral RNA quantitation was determined by RT-qPCR and expressed as number of SARS-CoV-2 E genome copies. Data represents the mean ± S.E.M of at least triplicate biological samples assayed in duplicate by RT-qPCR and is representative of two independent experiments. **d,** SARS-CoV-2 virus (10^4^ PFU) was pre-incubated with NHS (0-25%) or with 25% heat inactivated (HI) NHS (as control) for 1 hr at 37°C and then applied to Vero E6 TMPRSS2 cells. Infectivity determined after 3 days by plaque assay with readouts shown in PFU. Data are pooled from two independent experiments performed with duplicate biological samples. **b-d,** Each group compared to 0% NHS by one-way ANOVA with Dunnett’s multiple comparison test, **p < 0.01 and ***p < 0.001.

In addition to virolysis, we performed infectivity assays to determine whether NHS (sero-negative for SARS-CoV-2 spike antibody as determined by ELISA) had any impact on ability of SARS-CoV-2 to infect cells and produce infectious virus. Here, Vero cells were exposed to a fixed amount of SARS-CoV-2 that had been pretreated with NHS or heat-inactivated NHS (HI NHS). No loss of SARS-CoV-2 infectivity after exposure of virus to as much as 25% NHS was detected when cross-compared with exposure to HI NHS (Fig. 3d). Resistance to serum-mediated virolysis (Fig. 3b,c) was one possible explanation for how infectivity of SARS-CoV-2 was unimpeded in our experiment (Fig. 3d). We then went on to explore another possibility: that CL-11 binding could have had a compounding effect on viral infectivity.

### 4: Infectivity of SARS-CoV-2 is enhanced by bound CL-11

Certain human pathogens have adapted to complement allowing the microrganisms to escape destruction and promote tissue invasion^27–30^. We constructed an assay where respiratory epithelial cells were incubated with SARS-CoV-2 that had been coated or not with CL-11, followed by quantification of virus released into the culture supernatant using a plaque forming assay (PFA) in Vero cells. In an exploratory experiment we incubated SARS-CoV-2 (England 02/2020) with rCL-11 over a range of pre-determined concentrations up to 10µg/ml (Fig. 1a,b). Remarkably, infectivity of the CL-11-opsonised virus was up to 3.2-fold higher than for non-opsonised virus, based on the titre of virus released from infected bronchial epithelial BEAS-2B cells engineered to express ACE2 (Fig. 4a,c). We also observed that lung epithelial Calu-3 cells, which naturally express ACE2 and entry co-factors TMPRSS2 and TEMPRSS4^31^ exhibited a similar degree of susceptibility to rCL-11-treated virus as BEAS-2B (Fig 4b,c). Likewise, we also detected a 3-4 fold enhancement of infection with the highly transmissible and infectious B.1.1617.2 variant of SARS-CoV-2^32,33^ in BEAS-2B and Calu-3 cells respectively (Fig. 4 d-f). Generally, the amplifying effect of rCL-11 was dose-dependent with a peak effect between 1-10µg/ml (Fig. 4a-f). The reproducibility of the opsonising effect of rCL-11 for different cells and different viral strains is indicative of conserved virus-expressed glycans critical for CL-11 binding^34^.

**Fig. 4.**
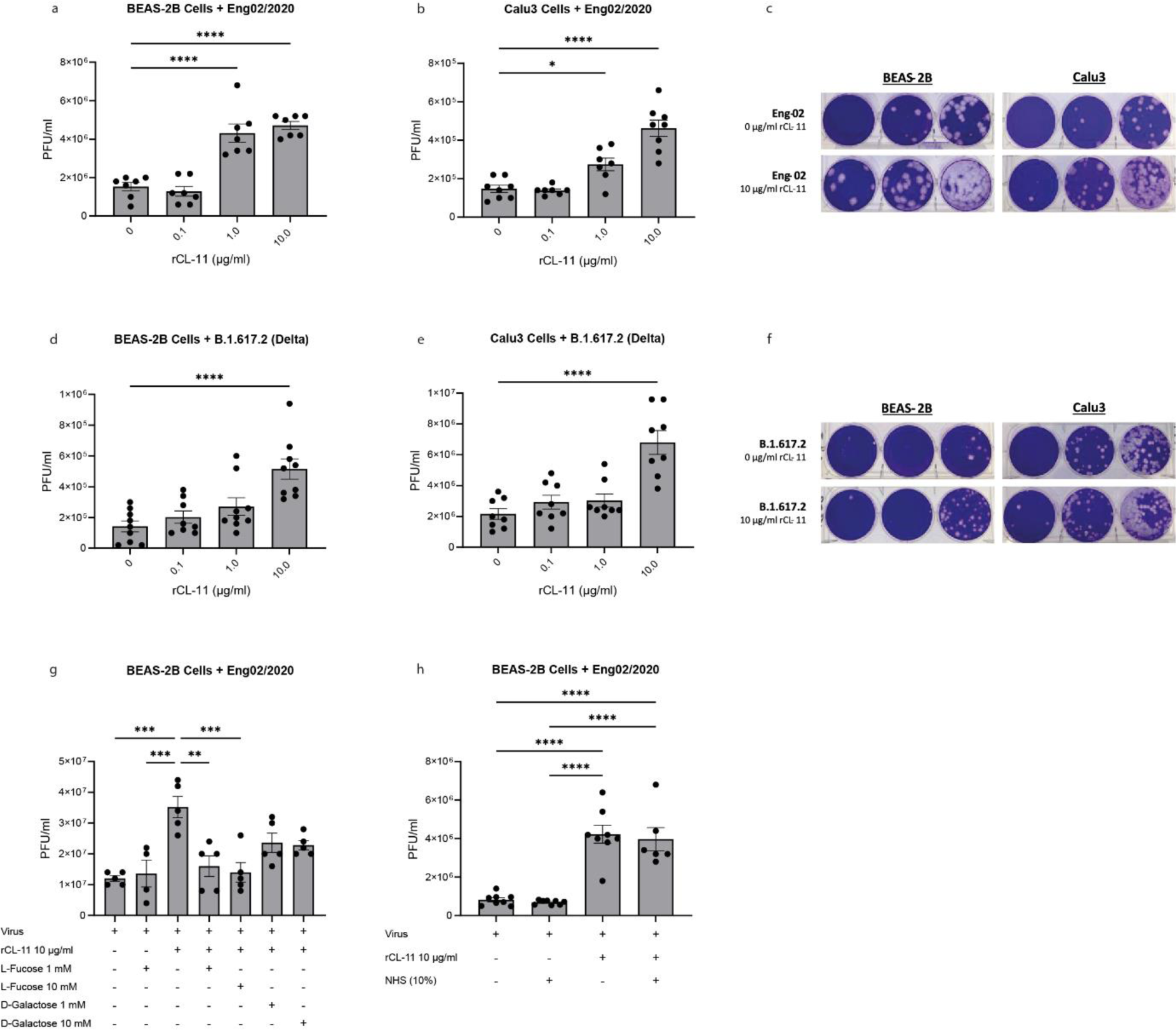
SARS-CoV-2 opsonised with CL-11 drives increased infectivity of respiratory epithelial cells. **a,d,** Enhancement of BEAS-2B cell infection with SARS-CoV-2 variants pre-treated with rCL-11 (0.1, 1.0 and 10.0 µg/ml) or media for 1 hr at 37°C. (**a**) England/02/2020 (MOI 0.005) and (**c**) B.1.17.2 (MOI 0.05). Virus titres determined in the supernatant after 24 hr by plaque assay in Vero E6 TMPRSS2 cells. Data are pooled from two (**a**) or three (**d**) independent experiments and represent the mean ± S.E.M of triplicate or quadruplicate biological samples. Each group compared to 0 µg/ml rCL-11 by one-way ANOVA with Dunnett’s multiple comparison test, * <0.05, and ***p < 0.001 **b, e,** Enhancement of Calu-3 cell infection with SARS-CoV-2 variants pre-treated with rCL-11(0.1, 1.0 and 10.0 µg/ml) or media for 1 hr at 37°C. (**b**) England/02/2020 (MOI 0.05) and (**d**) B.1.17.2 (MOI 0.05). Virus titres determined in the supernatant after 24 hr by plaque assay in Vero E6 TMPRSS2 cells. Data are pooled from 2 independent experiments and represent the mean ± S.E.M of quadruplicate biological samples. Each group compared to 0 µg/ml rCL-11 by one-way ANOVA with Dunnett’s multiple comparison test, ****p < 0.0001. **c,f**, Representative images of crystal violet stained plaque assay plates of BEAS-2B or Calu-3 cells infected with either England/02/2020 (c) or B.1.617.2 (f) or opsonised with 10mg/ml rCL-11. **g.** L-fucose inhibits the enhancement of SARS-CoV-2 infectivity driven by CL-11 in BEAS-2B cells. Quantification of SARS-CoV-2 (England/02/2020), as detailed above, in the supernatant of BEAS-2B cells 24 hr after infection with virus (MOI 0.05) pre-treated with media or with rCL-11 (10.0 µg/ml) in the presence or absence of L-fucose or D-galactose (1-or 10 mM final concentration). Data represents the mean ± S.E.M of five replicate biological samples and are representative of two independent experiments. *P* values were determined by one-way ANOVA with Tukey’s multiple comparison test. ** p<0.01 and ***p < 0.00. **h.** Infectivity of SARS-CoV-2 (England/02/2020) opsonised with CL-11 is not inhibited by NHS. Quantification of SARS-CoV-2 England/02/2020 (as detailed above) in the supernatant of BEAS-2B cells 24 hr after infection with virus (MOI 0.005) pre-treated with media or with rCL-11 (10.0 µg/ml) spiked with or without 10% NHS. Data are pooled from 2 independent experiments and represent the mean ± S.E.M of quadruplicate biological samples per experiment. *P* values were determined by one-way ANOVA with Tukey’s multiple comparison test. ****p < 0.0001.

To corroborate the importance of glycan recognition by CL-11 for enhanced infectivity of SARS-CoV-2, we preincubated rCL-11 with monosaccharide to saturate the carbohydrate-binding site on rCL-11 before mixing with the virus. We determined that preincubation of rCL-11 with L-fucose (1mM or 10mM) fully ablated the enhancing effect of CL-11 (Fig. 4g). D-galactose, which has lower binding avidity for CL-11 compared to L-fucose^7^ had a partial blocking effect (Fig. 4g). These experiments indicated that interaction via the CRD of CL-11, rather than the CLD, was primarily responsible for driving the enhanced infectivity of SARS-CoV-2.

The above infectivity experiments were conducted in the absence of exogenous complement. To understand whether the effect of CL-11 upon viral infectivity was maintained in the presence of complement, we performed assays in which rCL-11 had been spiked with NHS (10%) before mixing with SARS-CoV-2. To that end, we included adequate provision of complement factors (as indicated by Fig. 2) alongside rCL-11 to test the impact of complement factors on enhanced viral infectivity driven by rCL-11. We observed that CL-11-enhanced virus infectivity was not dependent on or limited by the provision of complement excess in this setting (Fig. 4h).

### 5: Respiratory epithelial cells infected by SARS-CoV-2 are sensitive to cell-autonomous complement

We envisage that locally produced CL-11 interacting with SARS-CoV-2 facilitates infectivity at the portal of entry and, in turn, virus-infected cells autoactivate complement through CL-11 binding. Suporting this hypothesis, we found that BEAS-2B cells infected with SARS-CoV-2 markedly upregulated gene expression of CL-11 and the core components of complement, C3 and C5. This occurred over time and in proportion to the virus dose (MOI), (Fig. 5a-c). Using confocal microscopy, detection of CL-11 and C3d proteins at the cell surface (of non-permeabilised cells) was most evident at higher virus MOIs (Fig. 6a,d, and Supplementary Fig 2 respectively), notably associated with the subpopulation of BEAS-2B cells expressing cell surface SARS-CoV-2 spike (spike^+^, Fig. 6c,f, respectively). Co-localisation of staining for cell-surface SARS-CoV-2 spike protein and CL-11 (Fig. 6d, top row) or C3d (Fig 6d, top row) was partial, meaning that the area of CL-11 and complement deposition was not limited to foci where spike protein was detected but it affected a wider surface area of virus-stressed cells. Complement activation proceeded to form membrane attack complex C5b-9 that was increased at the higher MOI of SARS-CoV-2, in spike^+^ expressing cells compared to uninfected cells (Fig. 5i). Collectively, these observations suggest that cell stress in general and SARS-CoV-2 spike protein at the cell surface in particular induced CL-11 binding and complement deposition following viral infection.

**Fig. 5.**
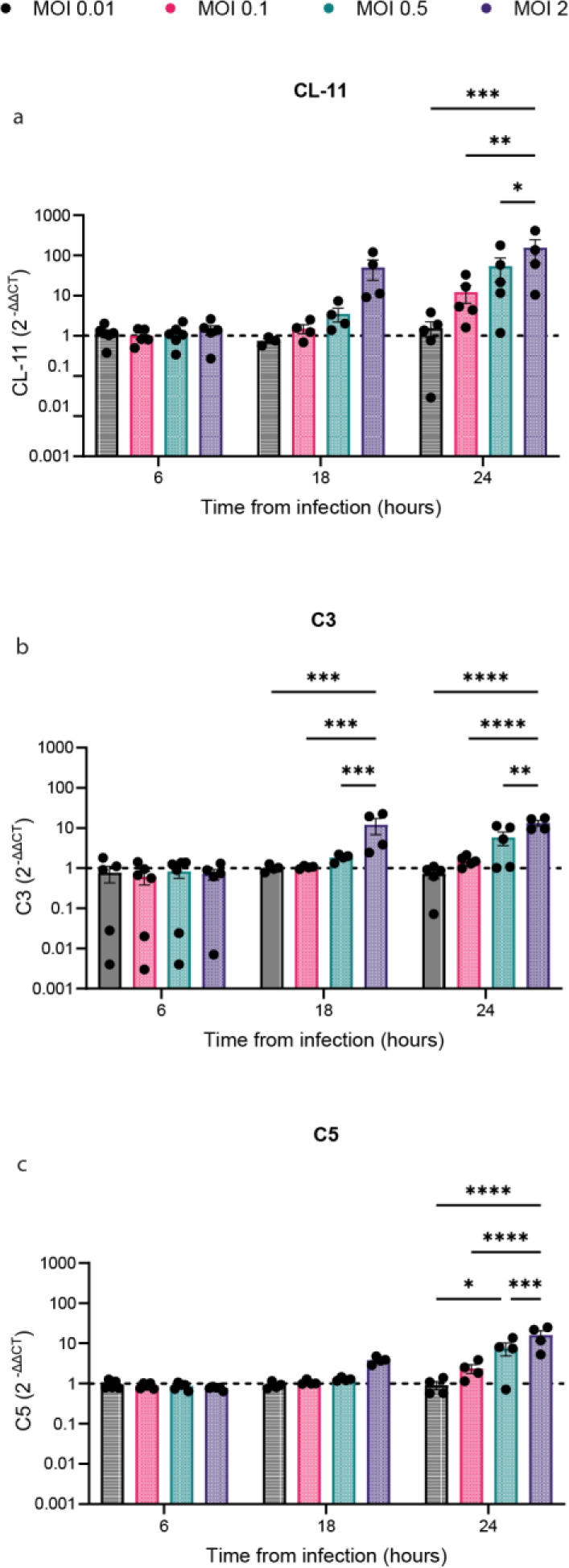
SARS-CoV-2 infection of bronchial epithelial BEAS-2B cells drives Cl-11 and complement gene expression. **a-c** Fold mRNA induction (2^−ΔΔCt^ method) of (**a**) CL-11, (**b**) C3 and (**c**) C5 (by qRT-PCR) after infection of BEAS-2B cells with SARS-CoV-2 (England/02/2020) over a range of MOIs (0.01, 0.1, 0.5 and 2.0) at 6, 18 and 24 h pi. Data represent mean ± S.E.M from n=4 independent experiments. Each group (within each time point) compared by two-way ANOVA with Tukey’s multiple comparison test. *p < 0.05, **p < 0.01, ***p < 0.001 and ****p < 0.0001.

**Fig. 6.**
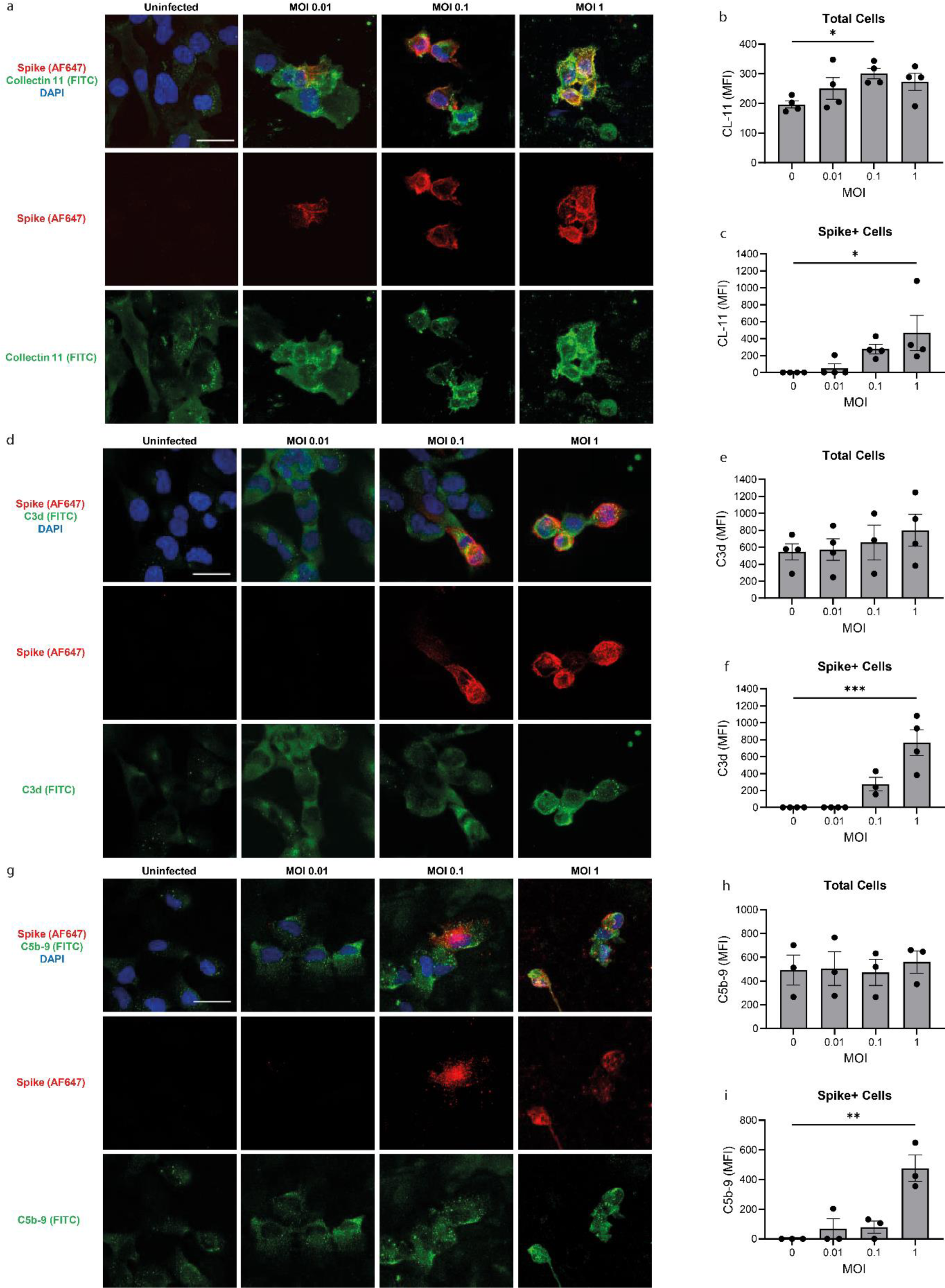
Bronchial epithelial BEAS-2B cells infected by SARS-CoV-2 are sensitive to cell-autonomous complement. **a, d, g,** Representative confocal microscopic images of BEAS-2B cells, mock, uninfected at 24 hr after plating and 24 hr after infection with SARS-COV-2 (England/ 02/2020) over a range of MOIs (0.01, 0.1 and 1.0) and stained for cell surface spike protein (red), DAPI (blue) and either (**a)** cell surface CL-11 (green), (**d)** cell surface C3d (green) or (**g**) cell surface C5b-9 (green). Original magnification: x 60, scale bars: 30 mm. Bar graphs show quantification of the MFI (mean ± S.E.M) from four independent experiments with a minimum of five images per condition per experiment for (**b**) CL-11 on all (total) cells imaged, (**c**) CL-11 on spike+ cells, (**e**) C3d deposited on all cells, (**f**) C3d deposited on spike+ cells, (**h**) C5b-9 on all cells and (**i)** C5b-9 on spike+ cells. Statistical comparisons are made by one-way ANOVA with Dunnett’s multiple comparisons between mock (uninfected) and SARS-CoV-2 infected conditions at each MOI (**e, f, h, j, k, l**). *p < 0.05, **p < 0.01 and ***p < 0.001.

## Discussion

Our data suggests CL-11 (a C-type lectin) is an important first-line defence of innate immunity produced at the respiratory epithelium at the point of contact with airborne pathogens such as SARS-CoV-2. Intriguingly, our study reveals that while production of CL-11 is enhanced by virus-infected respiratory epithelial cells and the lectin is directly bound by virus to trigger complement activation, the virus is resistant to complement-mediated lysis. Yet, once infected by SARS-CoV-2, respiratory epithelial cells expressing the spike protein are targeted by endogenous CL-11 and downstream complement activation products. These findings are consistent with reports on post-mortem analysis of lung tissue^14,15^ providing evidence of profound complement activation associated with SARS-CoV-2 antigen, indicative of overwhelming inflammation and viral persistence at the portal of entry.

During our study a report was published showing immobilised SARS-CoV-2 spike trimer could deposit CL-11 from normal serum^35^. Our data significantly extend these findings with new insights into SARS-CoV-2 pathogenesis, using rCL-11 to probe the interaction with immobilised virus and spike protein and with membrane-expressed spike protein. Notably, we demonstrated, the sugar- and EDTA-inhibitable nature of the interaction was typical of C-type lectin binding by the CRD of CL-11, validating the integrity of the recombinant lectin generated with an in-house mammalian (CHO) expression system. In contrast, another recently published study using a commercial wheat germ-derived rCL-11 did not detect interaction with solid-phase immobilised spike protein of SARS-CoV-2^36^. In our hands, comparison of CHO-expressed rCL-11 with the wheat-germ-derived rCL-11 (from the same commercial supplier as in the reported study) confirmed strong binding with our CHO-expressed rCL-11 but found negligible binding of the wheat germ-derived rCL-11 (Supplementary Fig. 3). Correct protein folding required for functional activity of CL-11 could have differed between the two recombinant proteins. Equally, lack of post-translational modifications (including glycosylation, hydroxylation of proline and lysine residues in the CLD and disulphide bond formation) inherent in the wheat germ system may have accounted for the differences in the CL-11 binding detected. The data presented here and in an independent report^35^, along with the predicted glycan interaction sites on the spike trimer^19^, robustly support a role for CL-11 interacting with SARS-CoV-2.

Following complement activation, susceptible C3-opsonised viruses are cleared by phagocytosis or, in the case of enveloped viruses, through lysis by MAC^3,37^. Several viral pathogens, however, have evolved resistance to the effects of complement. Among the mechanisms of resistance, viruses can express novel proteins or capture host proteins that inhibit C3 or C4 convertases or block subsequent MAC assembly. For example, the NS1 protein of Flaviviruses leads to the recruitment of several different host complement-regulatory proteins that either inhibit the classical or alternative pathways of complement activation or inhibit the formation of MAC^38^. Likewise Nipah virus encodes a protease activity capable of cleaving C3b^26^. Another evasion mechanism is mounted by HIV-1, which adsorbs human GPI-anchored complement-regulatory proteins for protection against C3 cleavage and MAC formation^24^. We propose that selective pressure imposed by complement in a zoonotic reservoir has caused SARS-CoV-2 to evolve a mechanism that confers relative resistance to elimination by complement. It is entirely feasible, for example, that exposure to the complement system in bats^39,40^ may be a factor in the evolution of SARS-CoV-2 before transmission to humans.

Early host defences at epithelial surfaces are recognised to impact pathogen spread and disease sequelae^41^. A notable finding of our data was the effect of rCL-11 enhancing viral infectivity. While other common pathogens (e.g., HIV-1 and *E. coli*) use bound C3b to enhance the infectivity of cells expressing complement receptors^3,42^, we found no evidence for such a mechanism in our experiments where virus incubated with CL-11 in the presence of 10% NHS did not enhance viral infectivity over that with CL-11 alone. Therefore, C3b mediated opsonisation of SARS-CoV-2 was non-essential for CL-11 driven enhancement of infectivity in our system. This implies other properties of CL-11 require consideration. For example, CL-11 is an oligomeric protein whose multivalency in conjunction with reactivity to both viral and mammalian glycan ligands may permit cross-linking of the virus with the epithelial cell surface. This could possibly increase internalisation or processing of the virus by cells expressing the ACE2 host-cell attachment receptor. The property to usurp CL-11 may be important for Wuhan-like viruses with lower affinity for ACE2^43,44^ to establish infection in respiratory epithelia that express lower levels of ACE2 compared to other tissues such as heart and kidney^45^. We suggest that complement-resistance combined with enhanced infectivity mediated by CL-11 may act as pathogenicity factors in the course of infection.

Cell-autonomous damage by complement is a feature of epithelial cell stress in the presence infectious and sterile (e.g., hypoxia) noxious stimuli^46,47^. Our data show enhanced deposition of CL-11 and complement products (C3d and C5b-9) on SARS-CoV-2 infected respiratory epithelial cells, both at sites where spike protein is detected and extending to loci where no spike protein was present. This implies CL-11 interacts with both stress-associated and spike-associated glycans along the surface of SARS-CoV-2 infected cells. Our data are consistent with a primary role for the lectin complement pathway triggered by CL-11 binding to the infected cells. A secondary effect of the alternative pathway adding to the deposition of complement on respiratory epithelial cells is not ruled out by our data. Indeed, the secretome of bronchial and alveolar epithelial cells contains factors D and B^48^, which are key components of the alternative pathway and are brought into play by the lectin-pathway enzyme MASP-3 acting on factor D^49,50^.

CL-11 belongs to a family of collectins that includes the structurally related MBL. While CL-11 and MBL share similarities including the range of glycans recognised, only CL-11 is synthesised in peripheral tissues, noticeably at epithelial surfaces including the lung^6,7^. An analysis of MBL has defined a potential role for neutralising SARS-CoV-2, relevant to the prevention of systemic spread of infection^36^. The difference here is that our study (of Cl-11) defines a potential role for CL-11 in the regional tissue response to the virus at the portal of entry. Future, *in vivo* studies may be best placed to analyse the biological significance of this compartmental model, which has implications for how, when and where treatment can be effectively delivered during early exposure to virus.

In conclusion, our results advance the notion that SARS-CoV-2 usurps the local defence system provided by CL-11 at the portal of viral invasion of the respiratory tract (Fig. 7). Virus treated with CL-11 was not only resistant to complement lysis, but the CL-11 opsonised virus displayed enhanced infectivity of respiratory epithelial cells. Additionally, SARS-CoV-2 infected cells demonstrated self-inflicted damage through CL-11 and complement deposition. In principle, this cycle of infection and cell injury can be broken by blocking CL-11 from interacting with glycan ligand, as shown by sugar blockade of the CRD of CL-11 in the present study, and previously in models of hypoxic tissue injury^50^.

**Fig. 7.**
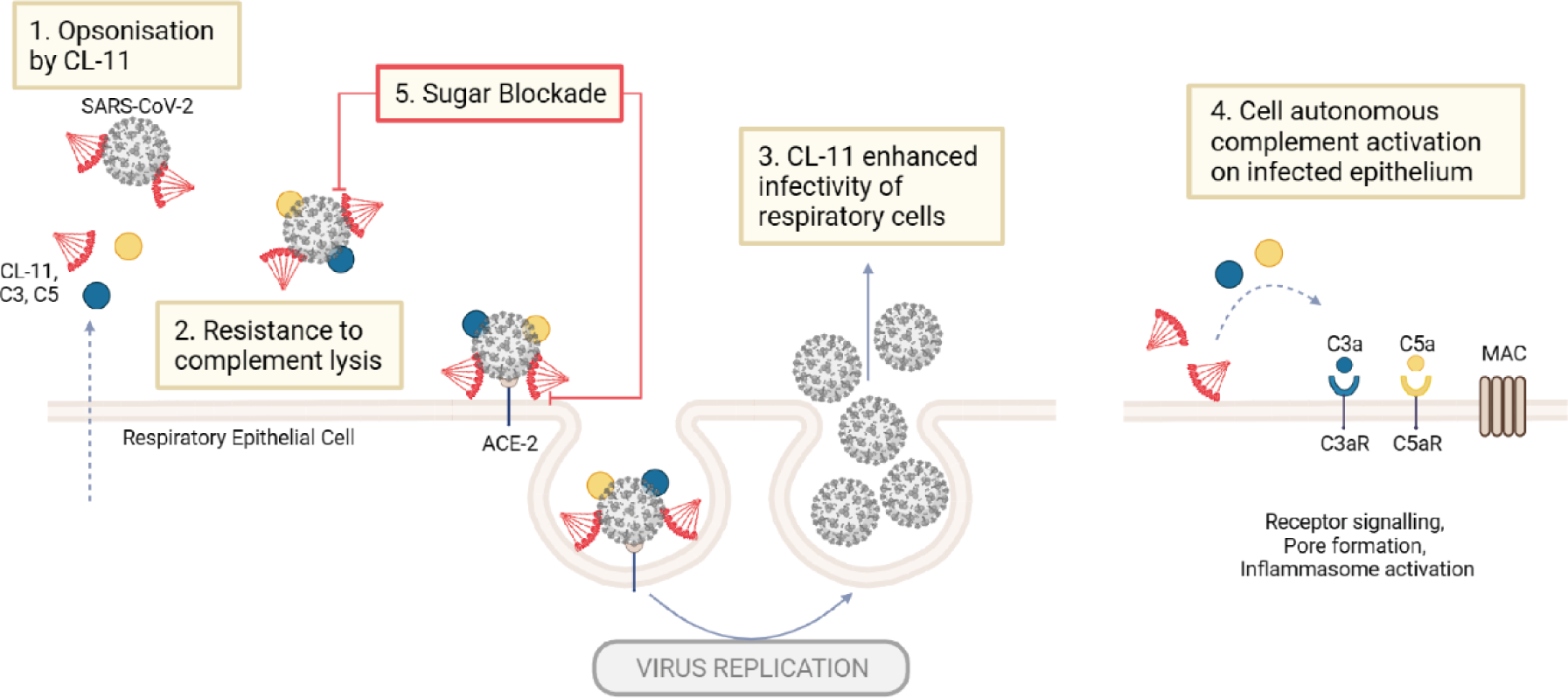
Schematic summarises the key findings of the study; demonstrating CL-11 modulation of respiratory epithelial cell invasion and injury by SARS-CoV-2. Epithelial cells of the respiratory tract are a source of CL-11, a secreted C-type protein interacting with glycan motifs on microbial and mammalian structures to stimulate complement activation. Our study demonstrates; (1) SARS-CoV-2 binds CL-11 and triggers complement activation to form membrane attack complex (MAC); (2) the virus however is resistant to lysis. (3); CL-11-opsonised virus exhibits enhanced infectivity, i.e., increased release of virus into the extracellular space of virus-infected cells by 3-4-fold; (4) virus-infected respiratory epithelial cells demonstrate cell-autonomous deposition of CL-11 and form complement products typically associated with inflammasome activation and membrane injury, (5), the cycle of enhanced infectivity can be interrupted by L-fucose given to saturate the carbohydrate-recognition domain of CL-11.

## Materials and Methods

### Recombinant proteins, serum and antibodies

Recombinant human collectin-11 (rCL-11) was expressed in mammalian CHO cells and purified as previously described^8^. A wheat-germ expressed rCL-11 (for comparative evaluation by ELISA) was purchased from Abnova. Unless specifically stated, CHO expressed rCL-11 was used in all experiments. All SARS-CoV-2 spike proteins were purchased from AcroBiosystems.

Normal human serum (NHS) pooled from male AB blood donors (pre-COVID) and confirmed SARS-CoV-2 spike seronegative by ELISA was obtained from NHS Blood and Transplant Services (NHBT). For some experiments NHS was heat inactivated (30 min at 56 °C) prior to use.

Full details of antibodies used are listed in supplementary information (Table 1).

### Cell lines and SARS-CoV-2 viruses

HEK293T (ATCC CRL-3216), Vero E6-TMPRSS2 (kind gift of Stuart Neil, King’s College London) and Calu-3 (ATCC HTB-55, kind gift of Jane McKeating, University of Oxford) were grown in Dulbecco’s modified Eagle’s medium (DMEM, GIBCO) supplemented with GlutaMAX, 10% fetal bovine serum (FBS) and 100 U/ml penicillin and 100 μg/ml streptomycin (Pen/Strep; Gibco). BEAS-2B-ACE2 (kind gift of Stuart Neil) were grown in RPMI 1614 (Gibco) supplemented with GlutaMAX, 10% FBS and Pen/Strep (Gibco) and maintained with 5 μg/ml blasticidin (InvivoGen) selection.

SARS-CoV-2 strain England 02/2020/407073 (referred to as England 02/2020) was obtained from Public Health England and strain B.1.617.2 (Delta variant) was a kind gift of Wendy Barclay, Imperial College London. Both isolates were propagated by infecting 60–70% confluent Vero E6-TMPRSS2 cells in T75 flasks. Supernatant was harvested 72 h post-infection after visible cytopathic effect, and filtered through a 0.22-μm filter (Sartorius) to eliminate debris, aliquoted and stored at −80 °C. Infectious virus titer was determined by plaque assay using Vero E6-TMPRSS2 cells. All work with infectious SARS-CoV-2 was carried out in a Containment Level 3 facility (Health and Safety Executive approvals, CBA1.295.20.1 and GM386/20.2).

### Inactivation of SARS-CoV-2 by UV irradiation

UV inactivation of virus was performed by exposure of SARS-CoV-2 (England 02/2020) at a maximal depth of 1.5mm in 6-well plates (Sarstedt) placed on ice, to a UV device (Jena Analytik) at 254 nm emission at a distance of 7.62cm for 5 min. All UV inactivated virus stocks were titrated by plaque assay to ensure no replicative virus remained.

### Plaque assay

Plaque assays were performed by infecting Vero-E6-TMPRSS2 cells with serial dilutions of SARS-CoV-2 for 1 h at 37°C. Subsequently, 2x overlay medium (DMEM with 2% FBS and 0.1% agarose) was added, and infected cells were fixed with 4% paraformaldehyde (PFA) at 72 h after infection and stained with crystal violet. Plaques were counted and virus titre determined.

### Binding of rCL-11 to immobilized SARS-CoV-2 proteins

Nunc MaxiSorp 96-well plates were coated with either UV-inactivated SARS-CoV-2 England/ 02/2020 or with SARS-CoV-2 spike protein, Wuhan or Omicron BA.1 (AcroBiosystems) over a concentration range in carbonate/bicarbonate buffer for 18h at 4°C. The plates were washed four times with Tris buffered saline (10mM Tris, 145mM NaCl, 0.05% Tween-20, 2mM CaCl2 pH7.4) before blocking with 0.1% bovine serum albumin (BSA) in Tris buffer for 1h at RT. Plates were then rewashed and incubated with rCL-11 (CHO cell produced) diluted in 10mM Tris, 145mM NaCl, 2mM CaCl2 for 1h at RT. Plates were then washed and incubated with rabbit anti-human CL-11 antibody (1:1000, Abcam) for 1h at RT. Plates were re-washed and incubated with goat anti-rabbit IgG-horseradish peroxidase (HRP) antibody (1:3000, Cell Signalling Technologies) for 1h at RT. After washing, plates were developed with 3,3’,5,5’-tetramethylbenzidine (TMB, Thermo Fisher Scientific) and absorbance read using a Hidex microplate reader (LabLogic) at 450nm. Values from blank wells were subtracted from sample wells. In certain experiments BSA (at the same molar concentration as rCL-11) was used as a control protein to rCL-11. The molecular mass of rCL-11 was 105 kDa corresponding to a trimer of polypeptides (based on SDS gels), the most abundant form of CL-11.

For the sugar blockade experiments, 96-well Nunc MaxiSorp plates were coated with SARS-CoV-2 Wuhan spike protein (AcroBiosystems) at 2µg/ml in carbonate/bicarbonate buffer for 18h at 4°C. Plates were then washed, blocked with 0.1% BSA (in Tris buffer) for 2h at RT and rewashed (as above). Plates were then incubated with rCL-11 (3μg/ml final concentration) and L-fucose or D-galactose over a titration range (0-200mM) or with 20mM EDTA, (all in 10mM Tris, 145mM NaCl, 2mM CaCl2) for 18h at 4 °C. Plates were then washed, incubated with rabbit anti-human CL-11 IgG (1:1000, Abcam) for 1h at RT, re-washed and incubated with goat anti-rabbit IgG-HRP antibody (1:3000, Cell Signalling Technologies) for 1h at RT and developed with TMB (as described above).

To compare the binding of rCL-11 produced in-house from transfected CHO cells^8^ with wheat-germ expressed rCL-11 (Abnova), the ELISA protocol described by^36^ was used. In brief, nickel high binding plates (Thermo Scientific, 15442) were coated with recombinant SARS-CoV2 spike protein (AcroBiosystems) at 2μg/ml in Tris buffered saline (10mM Tris base, 150mM NaCl2, 2mM CaCl2, pH 7.5) for 1h at 20°C. The plates were washed three times with Tris-Tween buffer (Tris buffered saline containing 0.1% Tween) and blocked with 2% BSA in Tris-Tween buffer for 2h at 37°C. Plates were then washed three times (as above), followed by addition of rCL-11 (CHO cell or wheat-germ expressed) at 0-9μg/ml in Tris buffered saline for 1hr at 37°C. After three washes with Tris-Tween buffer, a rabbit anti-human CL-11 IgG (1:10000, Abcam) was added to wells and incubated for 1h at 37°C. Plates were subsequently washed three times prior to addition of a goat anti-rabbit IgG-HRP antibody (1:3000, Cell Signalling Technologies), for 1h at RT. Plates were developed by addition of TMB, then quenched with 1M H2SO4 and the absorbance was read at 450nm as indicated above. Values from blank wells were subtracted from sample wells.

### Binding of rCL11 to cell surface expressed SARS-CoV-2 spike protein

HEK293T cells were transfected as previously described^51^ with either pcDNA3.1 (empty vector) or with SARS-CoV-2 Wuhan spike in pcDNA3.1 (kindly provided by Nigel Temperton, University of Kent). At 48 h post transfection, cells were harvested, washed in PBS containing 2% FBS and 1mM EDTA (FACS buffer) and 300,000 cells/sample were incubated with rCL-11 at 25µg/ml in PBS buffer containing 2mM CaCl2 or with the cognate buffer alone (as control), shaking for 1h at RT. In some experiments, rCL-11 was preincubated with either PBS, L-fucose (10 mM) or EDTA (10 mM) prior to incubation with spike or empty vector transfected HEK293T cells for 30 min at RT. The cells were then washed, stained with live/dead brilliant violet (BV510, Invitrogen) for 30 min at 4°C, washed, and stained with rabbit anti-human CL-11 IgG (20µg/ml, Abbexa) directly conjugated to Alexa fluor647(AF647, Invitrogen Alexa Fluor™ antibody labelling kit) for 45 min at 4°C. Cells were washed, stained with human SARS-CoV-2 anti-spike mAb (25µg/ml, P008_108)^52^ for 1h at 4°C followed by PE-rat anti-human IgG Fc (1:200, Biolegend) for 45 min at 4°C. Stained cells were washed, fixed with 4% paraformaldehyde (PFA) for 45 min at 4°C, rewashed, and acquired on a LSR Fortessa flow cytometer (BD). Data was analyzed using FlowJo software (version 10.1;TreeStar).

For the sugar blockade experiments rCL-11 was pre-treated with L-fucose (10mM) or EDTA (10mM) for 30 min at RT and then incubated with transfected HEK293T cells in PBS containing 2mM CaCl2 for 1h at RT shaking. Cells were stained and analysed by flow cytometry as indicated above.

### Complement activation assay

To measure C3a and C5a activation by SARS-CoV-2, 20μl NHS was incubated for 1 hr at 37°C with an equal volume of live SARS-CoV-2 (England/02/2020) from a 1:5 or 1:2 dilution series (as indicated) in PBS containing 2mM CaCl2. The reaction was stopped, and virus inactivated by incubation with PBS containing 1%-Tween-20 for 30 mins on ice. C3a-desArg and C5a levels were determined by ELISA (Hycult Biotech kit) according to the manufacturer’s instructions.

### Complement deposition assay

To measure activation and deposition of complement by SARS-CoV-2, Nunc MaxiSorp plates were coated with UV-inactivated virus (England/02/2020) in carbonate/bicarbonate buffer for 18h at 4°C. Plates were washed in TBS/Tween (10mM Tris, 145mM NaCl, 0.05% Tween-20, 2mM CaCl2, pH 7.4) and blocked with 3% BSA in TBS/Tween for 2h at RT. NHS diluted (0– 10%) in TBS plus 2mM CaCl2 was added to plates and incubated for 1h at 37°C. The plates were washed again and bound C3d or C5b-9 detected using a monoclonal anti-C3d antibody (1:500, Abcam) or a monoclonal anti-C5b-9 antibody against the neoepitope (1:100, Agilent Dako) and incubated for 1h at RT. Plates were subsequently washed, developed with TMB and reaction quenched with 1M H2SO4. The absorbance was read using a Hidex microplate reader (LabLogic) at 450nm. Values from blank wells were subtracted from sample wells.

### Viral lysis assay

Complement sufficient NHS (0-10% final concentration) was mixed 10^3^ PFU of SARS-CoV-2 and RNAse A (1mg/ml,Thermofisher) for 2 hr at 37°C. The reaction was stopped on ice and samples frozen 18 hr at −20°C to rupture damaged virions. Samples were then thawed, mixed with fresh RNAse A (1mg/ml, Thermo Fisher Scientific) and incubated for 1 hr at 37°C, followed by incubation with Proteinase K (1mg/ml, NEB) for 20 min at 37°C to remove RNAse A. Residual viral RNA was extracted and purified using the QiAMP Viral RNA extraction kit (QIAGEN) and quantified by RT-qPCR using a SARS-CoV-2 (2019-nCoV) CDC qPCR E primer and probe assay (Integrated DNA Technologies, #10006804). PCR reactions were performed using a QuantStudio-5 Real-Time PCR machine (Applied Biosystems) and analysed using QuantStudio Design and Analysis Software v1.5.2 (Applied Biosystems). Viral E genome equivalent copies were determined by standard curve, using an E gene standard kindly provided by Wendy Barclay, Imperial College London.

### Serum blocking infectivity assay

To test for serum-induced virolysis by an infection based assay, 10^4^ PFU of SARS-CoV-2 (England/02/2020) was incubated for 1 hr at 37 °C with an equal volume of pooled NHS (confirmed SARS-CoV-2 spike antibody negative by ELISA) at 0-25% (final NHS concentration) or with 25% HI NHS (56 °C for 30 min). Serial dilutions of NHS-virus mixtures were then titrated for infectious virus by plaque assay as stated above.

### Virus infections

BEAS-2B-ACE2 were seeded in 12-well plates at 2×10^5^ cells/ well in complete RPMI media for 24 hr and Calu-3 cells were seeded in 24-well plates at 2.5 ×10^5^ /well for 48 hr in complete DMEM. The media was then removed, cells washed with 2% HI FBS containing media and incubated for 1 hr at 37°C with mixtures of virus (equivalent to an MOI of 0.005 or 0.05 dependant upon the virus and cell line as indicated in the data figs) that had been preincubated for 1 hr at 37°C with rCl-11 at either 0, 0.1, 1.0 or 10.0 mg/ml (final concentration). The virus/rCl-11 mixture was then aspirated and cells washed in 2% HI FBS containing media. Cultures were incubated for 24 hr at 37°C in fresh 2% HI FBS media. Supernatants were then collected and stored at −80°C prior to infectious virus titre determined by plaque assay.

### RT-qPCR

RNA from infected and uninfected cells was extracted using a Qiagen RNeasy minikit (Qiagen; 74106) following the manufacturer’s instructions. Ten microliter of each extracted RNA was used to synthesise cDNA using a High-Capacity cDNA Reverse Transcription kit (Applied Biosystems, 4368813). RT-qPCR was performed using TaqMan Universal Master Mix (Applied Biosystems, 4304437) with specific TaqMan probes to quantify *COLEC11* (Hs00388161_m1), *C3* (Hs00163811_m1), *C5* (Hs01004342_m1) and *GAPDH* (Hs99999905_m1) all purchased from Applied Biosystems. RT-qPCR reactions were performed according to the manufacturers instructions using a QuantStudio-5 Real-Time PCR machine (Applied Biosystems) and QuantStudio Design and Analysis Software v1.5.2 (Applied Biosystems). Relative quantification was performed using the 2^−ΔΔCT^ method^53^ with sample data first normalized to GAPDH mRNA levels and then to values from uninfected cell control samples.

### Confocal microscopy

Coverslips in 24-well plates were pre-coated with poly-l-lysine for 18h at 4°C, then washed with PBS and seeded at a density of 2.5×10^4^ BEAS-2B cells per coverslip in 500ul complete RPMI media for 24 hr. Prior to infection, cells were washed with RPMI containing 2% HI FBS and then incubated with SARS-CoV-2 England/02/2020 (MOI of 0, 0.01, 0.1 and 1) for 1h at 37°C. Cells were then washed, replenished with RPMI containing 2% HI FBS and incubated for 24h at 37°C. Thereafter, cells were washed with PBS, fixed with 4% PFA for 30 min, washed again with PBS and then stored at 4°C until staining was performed. Fixed cells were blocked 30 min with PBS containing 20% normal sera (according to the species of the secondary antibody). For spike staining, PBS containing 1% BSA was used for blocking. Cells were then incubated with the primary antibody in blocking buffer, washed with PBS followed by an anti-species secondary antibody, in the same blocking buffer. After washing, cells were re-blocked prior incubation with the second primary antibody, followed by an anti-species second secondary antibody, followed by DAPI (Invitrogen) staining and mounting in PermaFluor (Epredia). The primary antibodies used were human mAb anti-spike (clone VA14_R37, 25μg/ml)^54^, rabbit anti-human C3d (1:200; Dako), and rabbit anti-human C5b-9 (1:200; abcam) all for 1h at RT, and rabbit anti-human CL-11 (1:100; Abbexa) 18h at 4°C. The secondary antibodies used were rat anti-human AlexaFluor-647 (1:200; Biolegend) and goat anti-rabbit FITC (1:200; Jackson Immunoresearch) for 1h at RT. Cells were imaged on a Nikon A1 inverted confocal (Eclipse Ti-E) microscope, using a Nikon Plan Apo lambda 60x/1.40 oil immersion lens, scanning at 1024 by 1024 pixels, resulting in a pixel size of 0.1 um/px. Images were acquired with a Galvano scanner controlled with NIS elements software. An average of 5 images were collected blind for each sample. All images were processed using Image J software (NIH) and quantitated as mean fluorescence intensity by circling the cell surface spike^+^ or spike^−^ cells as previously described^55^.

### Statistical analysis

Statistical analysis was performed using PRISM Software version 10.1 (GraphPad). Comparisons between multiple groups were performed using either a one-way or two-way ANOVA with multiple comparison tests as described in the figure legends. Data show the mean ± SEM, with significance shown on the figures and levels defined as *P<0.05, **P<0.01, ***P<0.001, ****P<0.00001.

### Data Availability

All relevant data are available from the authors upon reasonable request. A reporting summary for this Article is available as a Supplementary Information file.

## Acknowledgements

We thank Professors Wendy Barclay (Imperial College London), Jane McKeating (University of Oxford), and Stuart Neil (King’s College London) and Nigel Temperton (University of Kent) for the gift of invaluable cell lines and SARS-CoV-2 plasmids. We also thank Public Health England and Professor Wendy Barclay for SARS-CoV-2 isolates. We gratefully acknowledge the support of staff at the Nikon Imaging Centre, King’s College London, Dr Mark Howard for help with immunocytochemistry and Dr Giorgia Fanelli for help with flow cytometry. We likewise thank our colleagues for discussions and for their support. Schema figures were created with BioRender.com (free trial plan). This work was funded by the British Society for Antimicrobial Chemotherapy grant number BSAC-COVID-57 and funding from SEEK and supported by the National Institute of Health Research (NIHR) Biomedical Research Centre based at Guy’s and St Thomas’ NHS Foundation Trust and King’s College London. The views expressed are those of the authors and not necessarily those of the NHS, the NIHR or the Department of Health.

## Author Contributions

S.S conceived the hypothesis. L.S.K, A.P and S.S. co-designed the research; A.P, L.S.K, H.K., L.A.O’N, M.R, R.G, D.S, and T.W-G performed the research; K.J.D provided reagents, L.S.K, L.A.O’N, A.P, S.S, T.W-G and R.W analysed the data. A.P and T.W-G generated the figs. L.S.K, A.P and S.S and wrote manuscript. All authors contributed to editing the manuscript.

