## Supplementary File for "The complement pattern recognition molecule CL-11 promotes invasion and injury of respiratory epithelial cells by SARS-CoV-2"

### Supplementary Information

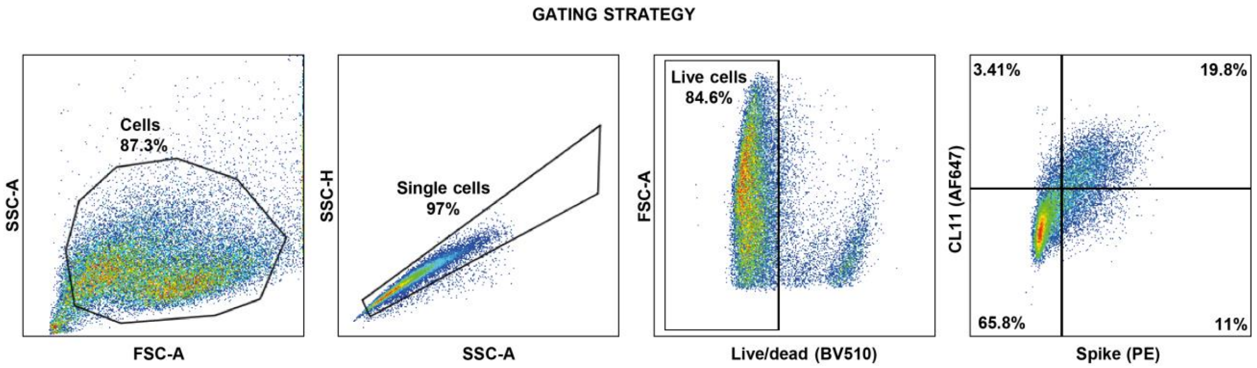


**Supplementary Figure 1. Gating strategy used to identify CL-11 on the surface of spike transfected HEK293T cells.**

Spike-transfected or empty vector transfected HEK293T cells were gated based on FSC-A and SSC-A. Doublets were excluded based on the SSC-A and FSC-A gate. Live cells were selected based on the live-dead (BV510) gate and were further examined for cell surface CL-11 and spike protein.

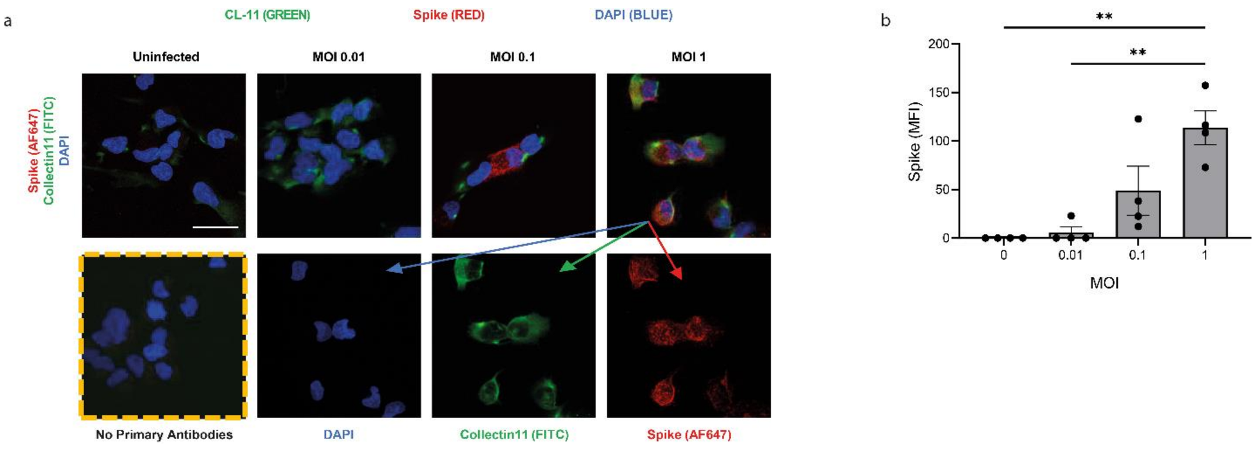


**Supplementary Figure 2. Efficiency of SARS-CoV-2 (England/02/20) infection of BEAS-2B cells.**

**a**) Upper panels; representative confocal microscopic images of BEAS-2B cells (mock uninfected) and 24 hr after infection with SARS-COV-2 (England/02/2020) over a range of MOIs (0.01, 0.1 and 1.0) and stained for cell surface spike protein (red, AF647), cell surface CL-11 (green, FITC) and DAPI (blue). Lower panels, represent (left to right), virus infected cells (MOI =1) with no primary antibodies (staining controls), followed by the merged image from upper row) MOI=1) split according to single colour immunofluorescence, red, AF647 (spike protein) and green, FITC (Cl-11) for comparison. Original magnification X60, Scale bars: 30 $\mu$m.

**b).** Bar graph shows quantification of cell surface spike protein MFI (mean ± S.E.M) from n=4 independent experiments with a minimum of n=4 images per condition (MOI) per experiment.

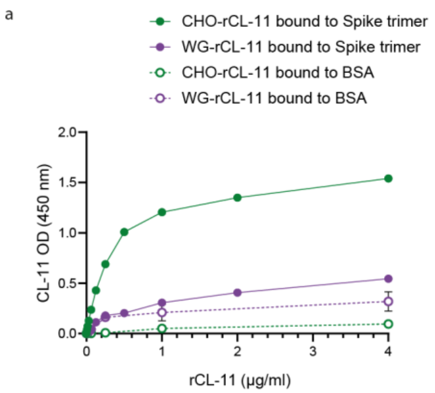


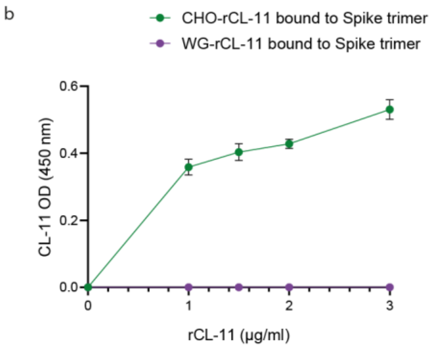


**Supplementary Figure 3. CHO-expressed but not wheat-germ-expressed rCL-11 binds to immobilised recombinant His tagged SARS-CoV-2 spike trimer protein.**

**a).** A fixed concentration of His-tagged SARS-CoV-2 spike trimer (2 μg /ml) or BSA (2 μg /ml) as a control was immobilised on Maxisorp ELISA plates and tested for capture of CHO-expressed recombinant CL-11 and wheat-germ expressed rCL-11 over a concentration range (0- 4 μg /ml) of each respective protein. Bound proteins were detected with rabbit anti-CL-11 antibody, followed by goat anti-rabbit HRP and TMB enzyme substrate. Binding measured by absorbance at 450 nm.

**b).** Nickel (high-binding ELISA plates) coated with His-tagged SARS-CoV-2 spike trimer (2 μg /ml) or BSA (2 μg /ml) as a control and tested for binding of CHO-expressed recombinant CL-11 and wheat-germ expressed rCL-11 according to the protocol described by Stravalaci et al (2022). Data (a, b) are enumerated by subtraction of values from blank wells from those recorded from sample wells and represent the mean ± S.E.M of n= 3 experiments performed in triplicate wells.

**Table 1: Key antibodies used**

| Antibody | Clone | Fluorochrome/ Conjugate | Source | Identifier | Application |
|---|---|---|---|---|---|
| Rabbit anti-human Collectin-11 | Polyclonal | - | Abbexa | abx003772 | Flow Cytometry ICC/Confocal |
| Rabbit anti-human Collectin-11 | Polyclonal | - | abcam | ab238585 | ELISA |
| Mouse anti-human C3d | 7C10 | - | abcam | ab17453 | ELISA |
| Rabbit anti-human C3d | Polyclonal | - | Dako/Agilent | A006302-2 | ICC/Confocal |
| Mouse anti-human C5b-9 | aE11 (neoepitope) | - | Dako/Agilent | M077701-8 | ELISA |
| Rabbit anti-human C5b-9 | Polyclonal | - | Abcam | ab55811 | ICC/Confocal |
| Anti-SARS-CoV-2 RBD | P008_108 | - | Doores lab | P008_108 | Flow Cytometry |
| Anti-SARS-CoV-2 RBD | VA14_R37 | - | Doores lab | VA14_R37 | ICC/Confocal |
| Rat anti-human IgG Fc | M1310G05 | PE | Biolegend | 410708 | Flow cytometry |
| Rat anti-human IgG Fc | M1310G05 | AF647 | Biolegend | 410714 | ICC/Confocal |
| Goat anti-mouse IgG (H+L) | Polyclonal | HRP | Jackson ImmunoResearch | 115-035-003 | ELISA |
| Goat anti-rabbit IgG (H+L) | Polyclonal | FITC | Jackson ImmunoResearch | 111-095-144 | ICC/Confocal |
| Goat anti-rabbit IgG (H+L) | Polyclonal | HRP | Cell signaling | 7074S | ELISA |
| Goat anti-rabbit IgG | Polyclonal | HRP | Cayman Chemical | 10004301 | ELISA |
